# T Cell Receptor Immunotherapy Drives Human Immunodeficiency Virus Evolution in Humanized Mice

**DOI:** 10.1101/574608

**Authors:** Alok V. Joglekar, Margaret Swift, Michael T. Leonard, John D. Jeppson, Salemiz Sandoval, David Baltimore

## Abstract

T cell receptor mediated immunotherapy using engineered Hematopoietic Stem/Progenitor Cells leads to durable partial suppression of HIV in humanized mice. Sustained viral suppression is accompanied by viral evolution under selection pressure. This study highlights the potential for TCR immunotherapy and the need to target multiple epitopes.

**Abstract:** Effective CD8+ T cell responses targeted to the KK10 epitope of HIV presented by HLA-B*27:05, a protective HLA allele, correlate with the ability to control infection without antiretroviral therapy (ART). Here, we report an immunotherapy approach using two B*27:05-KK10-specific T Cell Receptors (TCRs) isolated from HIV controllers. Immunocompromised mice engrafted with human Hematopoietic Stem/Progenitor Cells (HSPCs) encoding for the TCRs showed differentiation into functionally active engineered T cells. Following infection with HIV, both TCRs showed sustained, albeit modest, viral suppression over 32 weeks, accompanied by a concomitant increase in CD4+ T cells. Sequencing of viral quasi-species from the plasma of infected mice demonstrated clear evidence for viral evolution under selection pressure from the TCRs. The most commonly observed mutation in the KK10 epitope was L6M, which preserved viral fitness but showed attenuated recognition by the TCRs. These studies show that TCR-immunotherapy was able to suppress HIV infection long-term while driving HIV evolution in humanized mice.

## Introduction

HIV infection, if not treated with ART, can lead to uncontrolled viral replication and subsequent immune dysfunction. ART effectively suppresses viral replication, but requires strict, lifelong adherence to a treatment regimen. Interruptions in ART lead to viral rebound, underscoring that it is not a curative approach. A ‘functional cure’ may be achieved if viral replication is suppressed to manageable levels in the absence of ART (Katlama et al., 2013). A potential avenue for functional cure is to augment the immune system’s ability to combat HIV infection. This approach is fundamentally different from strategies that aim to make CD4+ T cells resistant to HIV infection. Augmenting the immune system through immunotherapy has the advantage of providing effector functions that may be able to suppress ongoing viral replication. One way to augment anti-HIV immunity is through enhancing HIV-specific CD8+ T cell responses. CD8+ T cells play an important role in controlling viral infections. Antiviral CD8+ T cells express antigen specific TCRs that recognize viral epitopes presented on class I HLA alleles on infected cells. Upon recognition of their cognate epitope, CD8+ T cells rapidly invoke their effector function, leading to specific killing of the target cell and secretion of cytokines such as IFNγ. In HIV infection, CD8+ T cells exert rapid but strong selection pressure on the virus, causing viral evolution (Goulder et al., 1997; Migueles and Connors, 2015). HIV can mutate the epitopes targeted by antiviral T cells (McMichael et al., 2010), which may contribute to viral rebound. Viral evolution under selection pressure from natural antiviral CD8+ T cell responses can drive emergence of escape mutations that can either render the virus unrecognizable to the CD8+ T cells, or force the immune system to adapt to them (Iglesias et al., 2011; Ladell et al., 2013). Nevertheless, the potency of CD8+ T cells in suppressing HIV infection can be manipulated therapeutically. Importantly for immunotherapy, CD8+ T cells can be engineered to express an exogenous antigen receptor to repurpose their effector function to a desired target (Johnson et al., 2006). Enhancement of CD8+ T cells by expressing a Chimeric Antigen Receptor (CAR) has been shown to effectively suppress virus in vitro and in vivo (Ali et al., 2016; Leibman et al., 2017; Zhen et al., 2017). However, CARs target the hypervariable envelope glycoprotein, and thus are susceptible to immune escape seen with antibody-based approaches (Dingens et al., 2019; Dingens et al., 2017). Engineering CD8+ T cells with TCRs allows targeting of more conserved regions of HIV, such as the *Gag* polyprotein. TCR-based approaches studied thus far have been targeted to the SL9 epitope presented by HLA-A*0201 (A2-SL9-TCR) (Kitchen et al., 2009; Kitchen et al., 2012; Varela-Rohena et al., 2008).

In addition to engineering T cells with CARs or TCRs, a long-term protective approach should provide a self-renewing source of engineered T cells. Therefore, several reports have explored the use of engineered HSPCs for immunotherapy. Human HSPCs are capable of long-term immune reconstitution in immunocompromised mice. Moreover, engineering HSPCs leads to in vivo differentiation into functionally active engineered T cells (Giannoni et al., 2013; Gschweng et al., 2014; Kitchen et al., 2009; Kitchen et al., 2012; Vatakis et al., 2011; Yang and Baltimore, 2005). The testing of HSPC-based approaches for TCR immunotherapy for HIV has been largely restricted to the SCID/hu or BLT mouse models that require human fetal tissue and thus are limited by availability of HLA-typed tissue (Carrillo et al., 2018) and regulatory hurdles. Therefore, research performed on TCR-immunotherapy approaches for HIV infection has been limited. Kitchen *et al* reported that HSPCs engineered with an A2-SL9-specific TCR differentiated into engineered T cells in SCID/hu mice, and showed viral suppression over 6 weeks. Due to weak selection pressure, long-term control of viremia was not observed. Furthermore, there was no evidence of viral evolution in these mice.

In this study, we tested an HSPC-based immunotherapy approach using TCRs specific to the KK10 epitope presented by HLA-B*27:05 (B27-KK10-TCRs). HLA-B*27:05 is a ‘protective’ HLA allele that is enriched among HIV controllers (Kiepiela et al., 2004). The CD8+ T cell responses in HLA-B*27:05+ patients are dominated by B27-KK10-specific T cells (Payne et al., 2010). Moreover, B27-KK10-specific CD8+ T cells are critical for HIV control through their superior functional capabilities (Buckheit et al., 2012a; Buckheit et al., 2012b; Chen et al., 2012). The properties of HLA-B*27:05 itself may affect thymic education of T cells, leading to more focused, and hence more protective, responses (Kosmrlj et al., 2010). The KK10 epitope is also hypothesized to be more structurally constrained as compared to the SL9 epitope (Schneidewind et al., 2008). Therefore, we tested if two B27-KK10-specific TCRs, EC27 and EC5.5, that we previously isolated from HIV controllers (Joglekar et al., 2018) could be used for HSPC-based immunotherapy. Immunocompromised mice were engrafted with CD34+ HSPCs enriched from G-CSF mobilized peripheral blood from HLA-B*27:05+ healthy donors. In mice engrafted with HSPCs engineered with EC27 or EC5.5, viral suppression was sustained without antiretroviral therapy. Furthermore, we show evidence of viral evolution under selection pressure from the therapeutic TCRs. These results have important implications for the development of T cell based immunotherapy approaches for HIV infection.

## Results and Discussion

### Engineered HSPCs engraft NSG mice and differentiate into T cells

Two B*27:05-KK10-specific TCRs, EC27 and EC5.5 (referred to as FC5.5 previously), isolated from HIV controllers were used for immunotherapy (Joglekar et al., 2018). The TCRs were cloned into lentiviral vectors and linked to a cell surface marker, LNGFR, using 2A peptide sequences. CD34+ HSPCs enriched from G-CSF-mobilized peripheral blood (mPB-HSPCs) were obtained from HLA-B*27:05+ healthy donors. Neonatal NOD/SCID/IL2Rγc^−/−^ (NSG) mice were irradiated and injected intrahepatically with EC27-, EC5.5-, or Mock-transduced HSPCs (hereinafter referred to as EC27-, EC5.5-, or Mock-mice, respectively). Peripheral blood from the mice was collected and analyzed by flow cytometry to measure human cell engraftment and differentiation (Fig S1A). Engraftment of human CD45+ cells in peripheral blood of the mice was evident by 1 month, and remained stable for 4 months. At 4 months post-engraftment, 55/56 mice showed greater than 0.2% hCD45+ cells among all CD45+ cells, with an average engraftment of 5-15% (Fig 1A). At 4 months post-engraftment, the human CD45+ cell fraction consisted of 20-29% hCD3+ and 55-60% hCD19+ cells (Fig 1B). Engineering HSPCs with EC27 or EC5.5 did not affect the development of hCD3+ and hCD19+ cells in vivo. To assess the presence of engineered T cells, peripheral blood was stained with anti-LNGFR antibody and B27-KK10-dextramer. Analysis of LNGFR+Dextramer+ cells revealed that a mean of 11% of hCD3+ cells in EC27- and EC5.5-mice expressed the desired TCR (Fig 1C). The fraction of LNGFR+Dextramer+ cells is similar to the range of B*27:05-KK10-specific CD8+ T cells observed in patients (Chen et al., 2012). These results indicate that humanized mice engrafted with engineered HSPCs could differentiate into hCD3+ T cells bearing the desired TCR. Establishment of humanized mice using engineered mPB-HSPCs is of particular significance. Potentially, TCRs restricted to any HLA allele may be tested in this system. Moreover, the degree of control over performing these studies is far greater than approaches utilizing BLT or SCID/hu mice because it allows for selection of donors based on HLA type and for establishing large cohorts based on cell availability. Therefore, this study establishes the framework for testing HSPC-based immunotherapy in humanized mice using TCRs targeting any given HLA allele(s).

**Fig 1.**
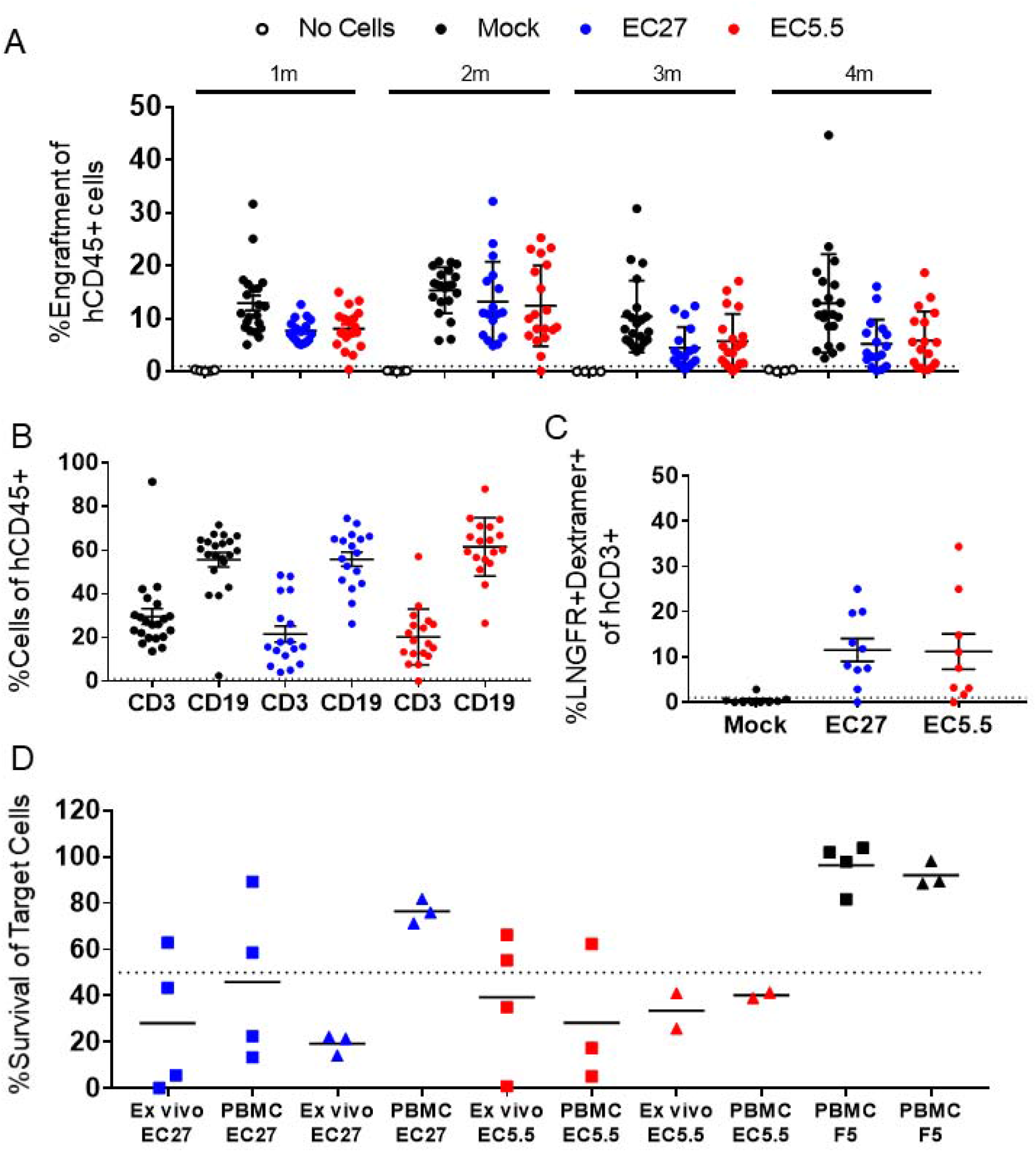
Differentiation of functionally active engineered T cells in humanized mice. A. Engraftment of hCD45+ cells in humanized mice over 1-4 months. %Engraftment is defined as %hCD45+ cell among total CD45+ cells. Each dot represents an individual mouse. Mean and S.D. are indicated with solid lines and error bars. Dotted line indicates 1% engraftment. B. Development of hCD3+ and hCD19+ cells at 4 months post-engraftment. Each dot represents an individual mouse. Mean and S.D. are indicated with solid lines and error bars. Dotted line indicates 1% of hCD45+ cells. C. Development of engineered T cells at 4 months post-engraftment. Each dot represents an individual mouse. Mean and S.D. are indicated with solid lines and error bars. Dotted line indicates 1% of hCD3+ cells. D. Measurement of ex vivo functional activity of engineered T cells from humanized mice. % Survival of target cells relative to Mock-effector cells was calculated. The source of effector cells, Ex vivo or from activated PBMCs, is indicated on the X-axis. For Ex vivo data: hCD45+ cells were isolated and expanded from engrafted mice, for PBMC data: hCD3+ cells from healthy donor PBMCs were expanded and transduced with EC27- or EC5.5-retroviral vectors. Each dot is one assay using effector cells pooled from 2-3 mice. Mean is indicated by the solid lines. Dotted line indicates 50% survival of target cells. F5 TCR was used as a control. Square dots indicate assays performed using KK10 peptide pulsed target cells, Triangular dots indicate assays performed using HIV-infected target cells.

### Human T cells from engrafted mice are functionally active

Upon confirming development of engineered T cells, ex vivo assays were performed to test their function. Previously described co-culture assays were used to measure KK10-specific cytotoxicity (Joglekar et al., 2018). GXR-B27 cells that express HLA-B*27:05 and express GFP upon HIV infection were used as target cells. Human CD45+ cells from spleens of engrafted mice were isolated by magnetic selection, activated and expanded ex vivo, and used as effector cells. GXR-B27 cells pulsed with KK10 peptide or infected with HIV^NL4-3^ were co-incubated with effector cells. Survival of target cells was measured at 24-hours after co-culture by flow cytometry (Fig S1B **and** C). Ex vivo expanded T cells from EC27- and EC5.5-mice showed cytotoxicity in response to KK10-pulsed target cells and HIV-infected target cells (Fig 1D). The effector function of hCD3+ T cells isolated from mice was comparable to primary T cells expressing EC27 and EC5.5 TCRs. These results show that engineered T cells developing from engineered HPSCs are functionally active against their target epitope.

### TCR Immunotherapy leads to durable but modest suppression of viremia

To test the ability of EC27 and EC5.5 to suppress HIV infection, two cohorts of humanized mice were generated. Human cell engraftment in peripheral blood of the mice was confirmed by flow cytometry. At 4-5 months post-engraftment, mice were challenged with 500ng p24 of HIV^NL4-3^ intraperitoneally. Peripheral blood was collected from the mice at regular intervals (every 4 weeks for cohort 22-23, and every 6 weeks for cohort 25-26). Frequency of human CD3+/CD4+ and CD3+/CD8+ cells was measured by flow cytometry (Fig S1D). Moreover, plasma from the mice was collected and used for viral load measurement by a clinical droplet digital PCR assay (Balazs et al., 2011; Strain et al., 2013). In both cohorts, EC27- and EC5.5-mice showed consistently lower viral loads than Mock-mice (Fig 2A and S2A). The frequency of hCD4+ T cells was significantly lower in challenged Mock-mice as compared to unchallenged mice, as expected. In both cohorts, frequency of hCD4+ cells was not significantly different among EC27-, EC5.5-, and Mock-mice (Fig 2B). At weeks 18-20 post-infection, a combined analysis of both cohorts showed 3.50-fold viral suppression by EC5.5 and 1.76-fold suppression by EC27. Viral suppression was accompanied by a 5.4% increase in %hCD4+ cells in EC5.5-mice. Importantly, at 18-20 weeks, 6 of 28 Mock-mice and 6 of 25 EC27-mice showed viral load below the level of detection, as compared to 12 of 23 EC5.5-mice (Fig 2C). Furthermore, the viral suppression observed in vivo was not donor-dependent, as similar results were seen in mice engrafted with cells from two different HSPC donors (Fig S2B). TCR-mediated viral suppression was observed at all time points, with the peak of 9.9-fold in EC5.5-mice, and 13.5-fold in EC27-mice. At the end of the study, at 32 weeks, EC27-mice showed 2.6-fold and EC5.5-mice showed 3.7-fold suppression (Fig 2D). Previous studies using ART in mice have shown over 10,000-fold suppression of virus (Olesen et al., 2016; Satheesan et al., 2018), whereas broadly neutralizing antibody administration has shown 30-100-fold suppression (Horwitz et al., 2013). However, in both cases, once the treatment was stopped, virus rebounded within 2-3 weeks to peak levels. In contrast, TCR immunotherapy using EC27 and EC5.5 showed viral suppression from the beginning of the study, and led to sustained suppression of virus over 32 weeks. However, it should be noted that the humanized mouse model used in this study does not reconstitute the immune system as robustly as the BLT or SCID/hu mouse models. Particularly, lower frequencies of antigen-presenting cells may impair T cell expansion and function, while reducing the sources of viral reservoirs. This is reflected in the fact that the mean viral loads observed in this study are 2-3-fold lower than those found in similar studies done with BLT mice. It is also possible that mice engrafted with HLA-B*2705+ HSPCs show inherently lower viral load. Previous results with BLT mice engrafted with HLA-B*57+ HSPCs showed ~3-fold lower viremia as compared to those engrafted with non-B*57+ HPSCs (Dudek et al., 2012). However, in our study, viral loads were compared to Mock-mice engrafted with HSPCs from the same donor. Overall, these results show clearly that TCR-immunotherapy suppressed viral infection long-term.

**Fig 2.**
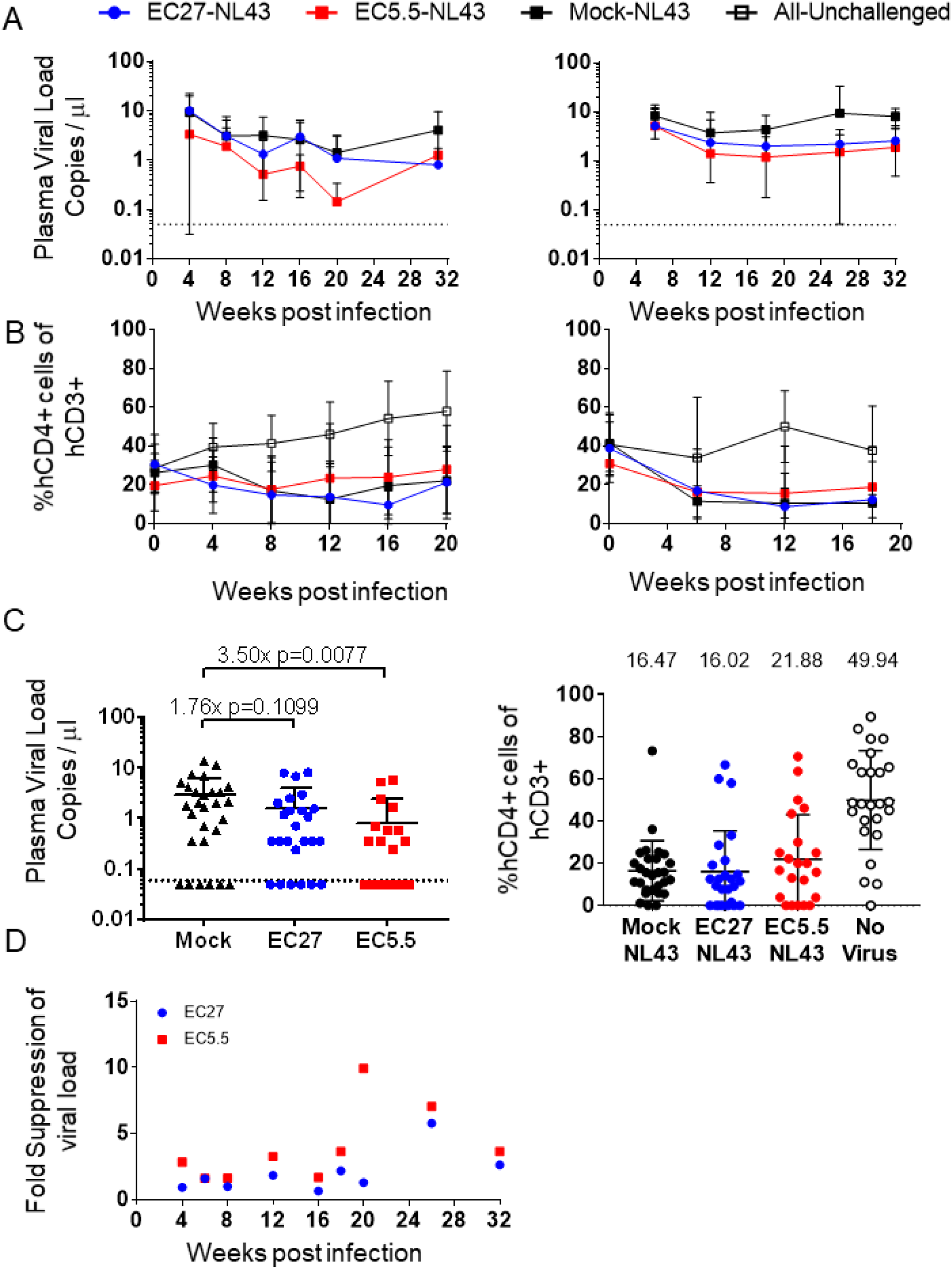
Long-term viral suppression by TCR immunotherapy. A. Viral load in the plasma from peripheral blood of HIV-challenged mice in cohort22-23 (left) and 25-26 (right). Dots and error bars indicate Mean and S.D. Dotted line indicates the limit of detection (0.05 copies per μl). B. Frequency of CD4+ T cells in peripheral blood of HIV-challenged mice in cohort22-23 (left) and 25-26 (right). Dots and error bars indicate Mean and S.D. C. Plasma viral load (left) and %CD4+ T cells at weeks 18-20 in both cohorts combined. Dots and error bars indicate Mean and S.D. Dotted line indicates the limit of detection (0.05 copies per μl). Left graph: Fold suppression of virus and p values from two-tailed Kolmogorov-Smirnov test as compared Mock-mice is shown above the data points. Right graph: Mean %CD4+ T cell frequency is shown above the data points. D. Fold suppression of viremia as compared to Mock-mice across both cohorts is shown.

### TCR-mediated viral suppression drives viral evolution in vivo

A likely scenario for TCR immunotherapy is immune escape due to viral evolution. To assess this possibility, viral quasi-species from the plasma of challenged mice were subjected to high throughput sequencing of the KK10 epitope. Predominant viral quasi-species constituting over 1% of the total diversity are depicted in Fig 3. At all 52 analyzed time-points, Mock-mice showed no evidence for viral evolution. In contrast, TCR-mice showed predominance of mutated KK10 epitope at 56 out of 64 measurements (35 of 39 for EC27 and 21 of 25 for EC5.5). Interestingly, at a majority of the time-points analyzed, the predominant mutation in the KK10 epitope was L6M, which changes KRWII**<u>L</u>**GLNK to KRWII**<u>M</u>**GLNK. Other mutations such as, R2K, R2G, R2Q, R2QL6M, R2KL6M, I5VL6M, L6I were also observed (Fig 3). An HLA-B*27:05-restricted control epitope, KY9, did not show detectable mutation in any of the mice (Fig S3). These results indicate clearly that ongoing selection pressure mediated by EC27 and EC5.5 TCRs is driving viral evolution. Interestingly, viral evolution was evident by 6 weeks post-infection; and in multiple instances, complete replacement of the predominant viral species occurred within 4 weeks. These results show maintenance of ongoing selection pressure.

**Fig 3:**
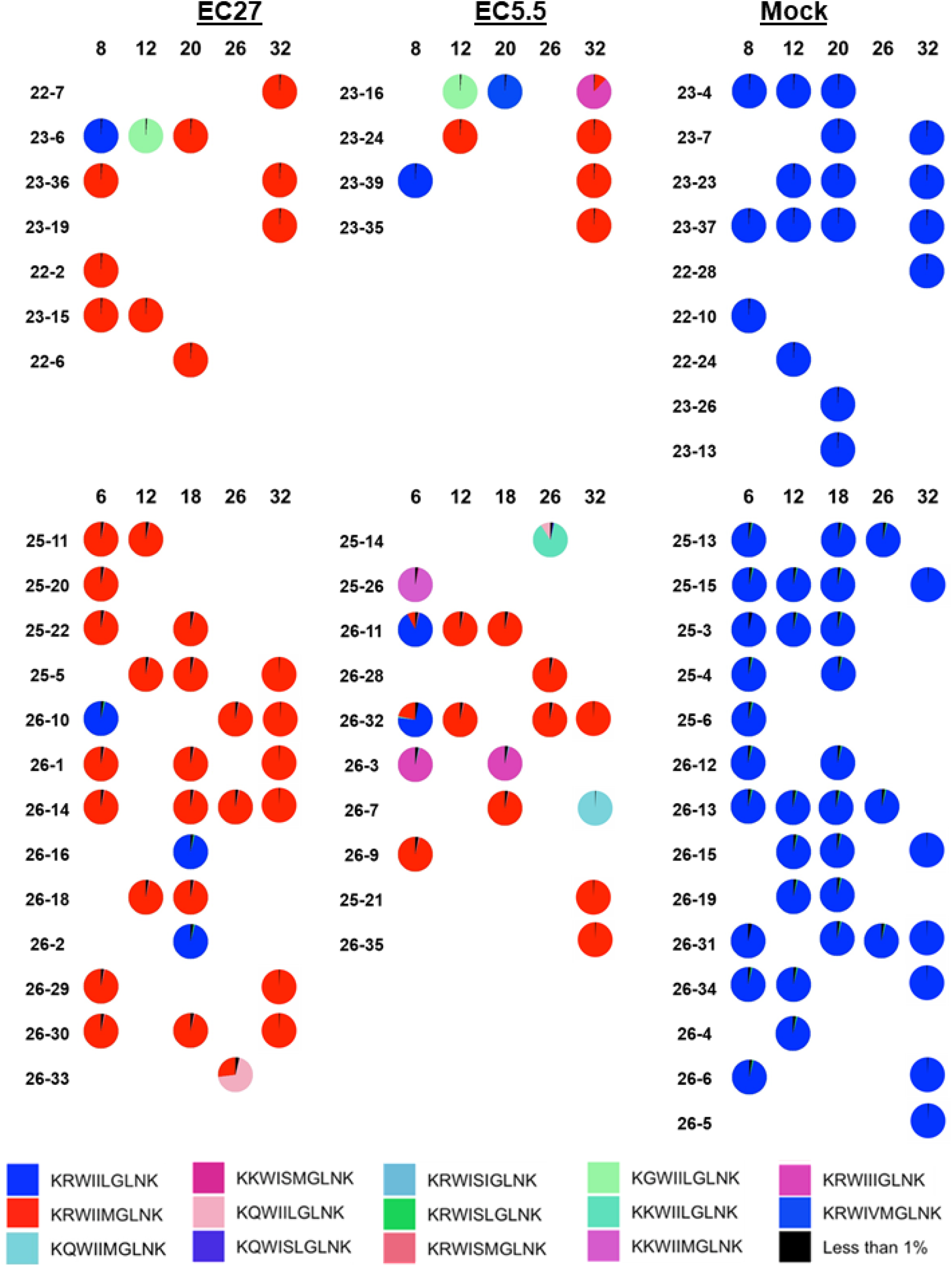
Evolution at the KK10 epitope in viral quasi-species in infected mice. Each pie chart indicates all viral quasi-species encompassing over 1% of the total sequence diversity at the KK10 epitope. For each TCR, mouse IDs are shown on the left and the week post-infection at which the sample was obtained is indicated on top. The top part of the figure shows data from cohort22-23, the bottom part shows data from cohort 25-26. The legend for the individual colors and their corresponding mutant epitopes is at the bottom. The wildtype epitope is represented by blue and the L6M epitope is represented by red.

### L6M mutation attenuates immune recognition without affecting viral fitness

The L6M mutation has been extensively documented in HLA-B*27:05+ patients as one of the most commonly found mutations in the KK10 epitope (Ammaranond et al., 2011; Lichterfeld et al., 2007; Lissina et al., 2014; Schneidewind et al., 2008). L6M mutation has little impact on viral fitness (Schneidewind et al., 2008; Schneidewind et al., 2007), which was confirmed by measuring infectivity of HIV^NL4-3^ encoding either the wildtype KK10 epitope or L6M mutation (Fig 4A). L6M is usually not associated with immune escape (Ammaranond et al., 2011; Streeck et al., 2008). However, some studies have shown reduced recognition of L6M by KK10-specific T cells (Chen et al., 2012; Iglesias et al., 2011; Xia et al., 2014). In our previous study, L6M mutation did not affect cytotoxicity when target cells were pulsed with saturating amounts of the peptide (Joglekar et al., 2018). Cytotoxicity assays using GXR-B27 cells infected with wildtype or L6M mutant viruses showed attenuated recognition of L6M (Fig 4B). Measurement of antigen sensitivity towards L6M indicated a 100-fold attenuation of recognition of L6M (Fig 4C). However, it must be noted that recognition of the KK10 epitope was not completely abolished by L6M, and thus it is a partial escape mutation. Predominance of the L6M mutation in infected mice can predict two potential scenarios. On one hand, mutating the KK10 epitope to L6M may represent a balance between viral fitness and immune selection pressure. This may be a result of attenuated recognition of L6M by the TCRs. However, sustained modest suppression of viremia would indicate ongoing selection pressure. Therefore, we posit that these mice represent a ‘viremic control’ scenario where the equilibrium between viral control and replication is maintained. On the other hand, L6M mutation is often compensatory of mutations at the R2 position. Previous studies have indicated that mutations such as R2G and R2K that impair immune recognition also impair viral fitness. The addition of L6M to these mutations improves viral fitness while maintaining attenuated viral recognition, leading to immune escape (Schneidewind et al., 2008; Schneidewind et al., 2007). It is conceivable that in the current study, emergence of L6M mutation may represent the first step towards emergence of true escape mutations. Replacement of L6M by R2QL6M was observed in only one mouse. In this case, our results would indicate a prolonged asymptomatic phase of infection after lowering the viral set point. However, as a period of 4-6 weeks is sufficient for complete replacement of the dominant species, we would expect emergence of escape mutations by 32 weeks more frequently. In addition, evidence for predominance of L6M mutation in long-term non-progressors (Ammaranond et al., 2011) also supports a ‘viremic control’ scenario.

**Fig 4:**
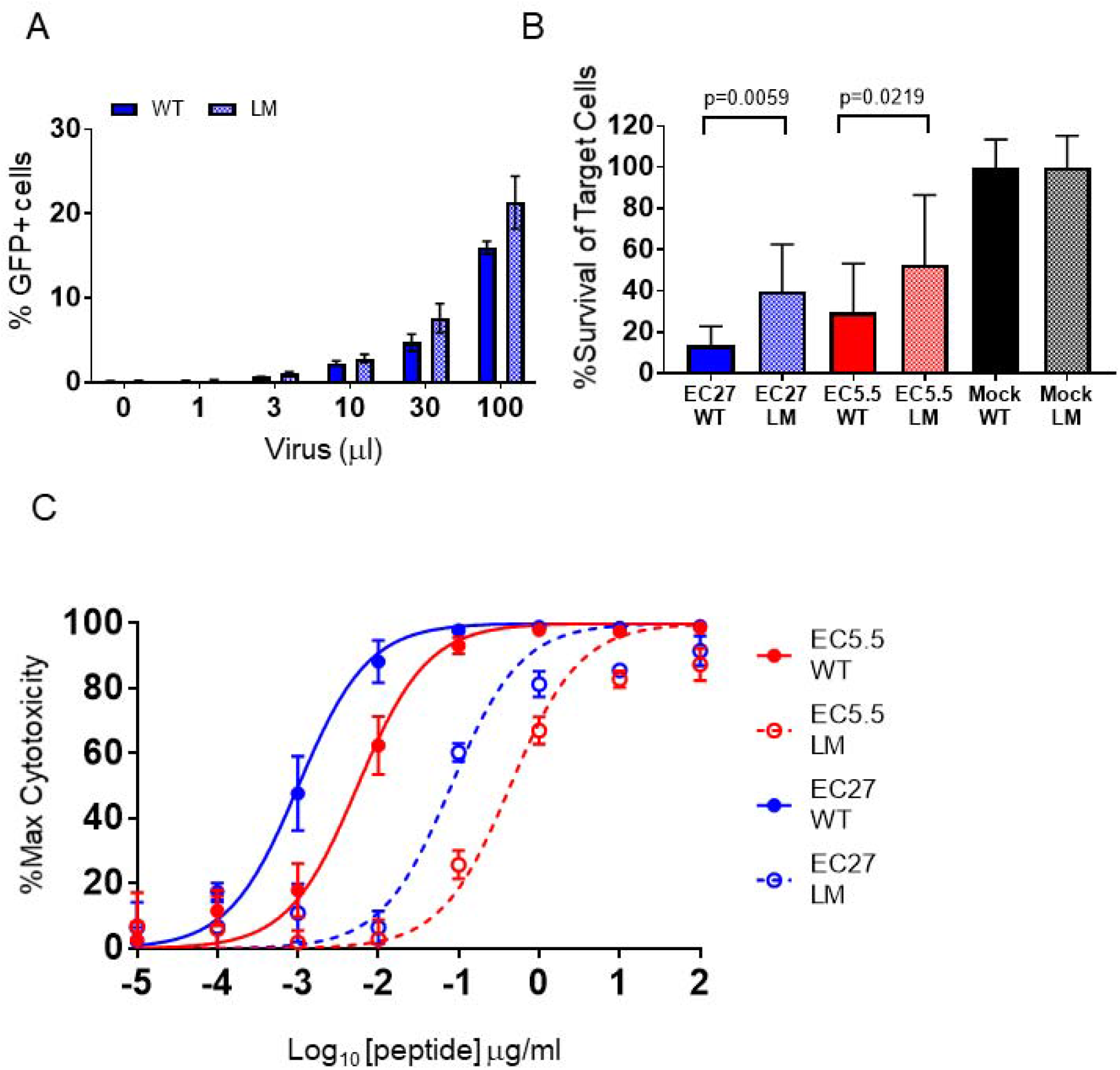
Effect of the L6M mutation on viral fitness and recognition by TCRs. A. Infectivity of the wildtype (WT) and L6M mutant (LM) virus on GXR-B27 cells. %GFP+ cells at 3 days post-infection are indicated. X-axis indicates the amount of virus used for infection. Bars and error bars indicate Mean and S.D. from n=3 biological replicates. B. Cytotoxicity towards GXR-B27 cells infected with WT or LM viruses. %Survival of target cells at 48 hours is indicated. Bars and error bars indicate Mean and S.D. from n=16 biological replicates. p values computed for One-way ANOVA with Sidak correction for multiple comparisons between WT and LM for each TCR are indicated. C. Antigen sensitivity towards WT and LM for both TCR. Cytotoxicity as a function of the amount of peptide used to pulse GXR-B27 cells is indicated. Non-linear regression fit curves are indicated by solid and dashed lines. Each point indicates Mean and S.D. from n=4 biological replicates.

## Concluding Remarks

In this study, we demonstrated sustained suppression of HIV mediated by TCR immunotherapy in humanized mice. Remarkably, 2.5-3.5-fold reduction in viremia was achieved at 32 weeks. These results highlight the efficacy of TCR immunotherapy for controlling viremia. To our knowledge, this is the first report of long-term suppression HIV achieved by TCR immunotherapy. These results pave the way for further development of this approach for more effective, long-term treatments with the eventual goal of achieving a functional cure. We also showed clear evidence for ongoing TCR-mediated selection pressure. Interestingly, most of the emergent viral quasi-species did not harbor true escape mutations in the KK10 epitope, indicating that they are still under immune pressure by the TCRs. Of note, a large majority of the mice harbored the L6M mutation, which is one of the most well-documented mutations in HLA-B*27:05+ patients. However, the occurrence viral evolution underscores that broadening of immunotherapy to target multiple epitopes may be necessary to enhance viral suppression.

There are caveats to this study which need to be considered. Firstly, the humanized mouse model used in this study does not reconstitute the immune system as robustly as the BLT or SCID/hu mouse models. Particularly, lower frequencies of antigen-presenting cells may impair T cell expansion and function, while reducing the cells that maintain viral reservoirs. Moreover, T cell function may also be impaired due to lack of HLA-matched thymic epithelium. However, our results show that T cell function is retained in ex vivo assays, alleviating this concern.

In summary, our results suggest that HSPC-based TCR immunotherapy is effective at long-term viral suppression despite driving viral evolution, and provide a basis for future testing of this approach in human clinical studies.

## Materials and Methods

### Primary cells and cell lines

GXR-B27 cells were a kind gift from Dr. Bruce D. Walker’s laboratory at the Ragon Institute of MGH, MIT, and Harvard. Jurkat (TIB-152) cells and HEK-293T (CRL-3216) cells were obtained from American Type Culture Collection (ATCC, Manassas, VA). CD34+ HSPCs from healthy donors were obtained from the Core Center for Excellence in Hematology at the Fred Hutchinson Cancer Research Center. Cryopreserved, magnetically enriched CD34+ HPSCs from G-CSF-mobilized leukapheresis product were obtained from two HLA-typed healthy donors. A low-resolution in-house PCR based assay, followed by a clinical 4-digit HLA typing assay was used to confirm that the donors were HLA-B*27:05+. Peripheral blood mononuclear cells (PBMCs) from healthy donors were obtained from the Virology core at the Center for AIDS Research at the University of California, Los Angeles.

### Cell culture conditions

Jurkat cells, GXR-B27 cells and PBMCs were cultured in RPMI1640 (Corning; 10-040-CV) supplemented with 10% FBS (Corning; 35-025-CV) and Pen/Strep (Corning; 30-002-CI) (R10). HEK-293T cells cultured in DMEM (Corning; 10-013-CV) supplemented with 10% FBS and Pen/Strep (D10). HSPCs were thawed in IMDM (Gibco; 12440-053) supplemented with 20% FBS and Pen/Strep (I20), and cultured in X-VIVO15 (Lonza; 04-744Q) supplemented with Pen/Strep (0XTD). All of the cells were cultured in humidified incubators at 37 °C with 5% CO2 (VWR).

### HSPC transduction and culture

Cryopreserved HSPCs were thawed in I20, centrifuged at 500g for 10 min at 4 °C and resuspended in 0XTD supplemented with 100 ng/ml each of Stem Cell Factor (SCF, Miltenyi Biotec, 130-096-695), Flt3 Ligand (Flt3L, Miltenyi Biotec, 130-096-479), and Thrombopoeitin (TPO, Miltenyi Biotec, 130-094-013), and plated on Retronectin-coated (Takara Bio; T100B) 12-well non-treated plates (Falcon; 351143). The cells were plated at a density of 3-6×10^5^ cells per well in 250 μL for 18 hours. Lentiviral vector was added to the cells at 1×10^8^ TU/ml to the cells. After 24 hours, the cells were harvested from the wells, pooled according to the vector used, and centrifuged at 500g for 10 min at 4 °C. Cell pellets were washed once with DPBS (Corning; 21-031-CV), and centrifuged at 500g for 10 min at 4 °C. Cells were resuspended in 0XTD at 10×10^6^ cells/ml and kept on ice until further use.

### Primary T cell transduction and culture

Cryopreserved PBMCs were thawed in R10, and cultured in R10 supplemented with 40 U/ml rhIL2 (Miltenyi Biotec; 130-097-746) and Immunocult anti-CD3/28 activator (Stemcell Technologies; 10971) for 4 days. Cells were then harvested, and transduced with MSCV-based retroviral vectors expressing EC27, EC5.5, or F5 TCRs. RD114-pseudotyped retroviral vectors expressing TCRs were packaged as described previously (Joglekar et al., 2019). Briefly, HEK-293T cells were plated on poly-L-Lysine-coated (Sigma; P4707) 10 cm plates (Corning; 430167) at 5×10^6^ cells per plate in D10. The cells were transfected 24 hours later with pMSCV-TCR, pHIT60, and pRD114. The medium was harvested 72 hours later, filtered through 0.45 micron filters (Millex-HV; SLHV033RS), and stored at −80 °C until use. For transduction, activated T cells were plated on Retronectin-coated 12-well non-treated plates, mixed with thawed viral supernatant, and spin-infected by centrifugation at 1111g for 2 hours at room temperature.

### Lentiviral vector packaging and titer determination

Lentiviral vectors expressing EC27 and EC5.5 TCRs along with LNGFR were constructed, packaged, and concentrated as described previously (Joglekar et al., 2018). Briefly, HEK-293T cells were plated on poly-L-Lysine-coated 15 cm plates (Corning; 430599) at 15×10^6^ cells per plate. The cells were transfected 24 hours later with pCCLc-MND-EC27 or pCCLc-MND-EC5.5 plasmid, pCMV-RΔ8.9, and pMDG-VSVG. Transfection medium was replaced 16 hours later with D10 supplemented with 10 mM Sodium Butyrate (Alfa Aesar, AAA11079-22) and 20 mM HEPES (Corning: 25-060-CI). Following butyrate induction, cells were washed once with DPBS and the medium was replaced with Ultraculture (Lonza; 12-725F) supplemented with Pen/Strep. The culture medium was harvested 48 hours later, filtered through 0.45 micron filters, and concentrated using Amicon Ultra-15 Filters (Millipore; UFC910096). Concentrated virus was stored at −80 °C until use. The titer was determined by transducing Jurkat cells and measuring vector copy number per cell using qPCR as described previously (Joglekar et al., 2018).

### Engraftment of NSG mice

NOD/SCID/IL2Rγc^−/−^ (NSG, NOD.Cg-Prkdc^scid^ Il2rg^tm1Wjl^/SzJ, The Jackson Laboratory, 005557) mice were purchased and bred in house. To obtain sufficient cohorts of neonatal mice, timed matings were setup with breeding trios. Upon confirmation of pregnancy, mothers were housed individually in cages. Neonatal, 3-5 day old pups were used for engraftment. Pups were first separated from the mother, transferred into sterile plastic containers along with bedding from the cage, and transported to the Irradiator. The pups were then irradiated at 100 cGy using a ^137^Cs radiation source, and transported back to the animal facility. Immediately following irradiation, the pups were injected intrahepatically with 5-7×10^5^ cells using an insulin syringe. To identify the experimental arm, the pups were tattooed on their paws with blue ink (Ketchum Inc). To avoid any technical bias, injections of EC27-, EC5.5-, or Mock-HSPCs were done on a rotating basis. After tattooing and injection, the pups were transferred back to their cage with their mother, along with the bedding. The pups were weaned from the mother at 3 weeks of life.

### Animal Identification and peripheral blood collection

At 1-month post-engraftment (or 4 months for cohorts 22-23 and 25-26), mice were earpunched to indicate their ID number. Mouse ID numbers were not correlated with blue tattoo indicating the TCR arm. At indicated time-points, mice were anesthetized under isofluorane (Piramal Healthcare), and retro-orbital blood collection was performed. Blood was collected using plain hematocrit tubes (Drummond; 1-000-7500-C/50) into K2-EDTA tubes (Terumo CapiJect; T-MQK or Greiner; 450480) and processed the same day. The blood collection tubes indicated only the animal IDs, and hence were processed in a blinded fashion. Plasma was collected where indicated and stored at −20 °C until further use.

### Tissue analysis

For collecting and analyzing tissues, mice were transported in cardboard containers to the laboratory and euthanized using carbon dioxide-mediated asphyxiation. Peripheral blood was collected by cardiac puncture using a 1 ml syringe attached to a 20G needle. The mice were then dissected under sterile conditions and spleens, thymi, and lymph nodes were collected. The tissues were mashed using the plunger of a 10 ml syringe on a 40 micron nylon filter (VWR; 10199-654). The filters were washed with MACS buffer (DPBS with 2% FBS) to collect cells. For bone marrow collection, tibiae and fibulae from one of the legs were collected and flushed with MACS buffer using a 27G needle through a 40 micron filter. For all tissues, staining for flow cytometry was performed as described below.

### Flow cytometry on mouse blood and tissues

Flow cytometry was performed using previously described protocol (Joglekar et al., 2018). Blood cell pellet or tissue cell pellet was resuspended in dextramer diluted in MACS buffer and incubated at room temperature for 20 min. Antibody mixture diluted in MACS buffer was added to the reaction followed by 20 min incubation at room temperature. The staining reaction was then subjected to RBC Lysis/Fixation (Biolegend; 422401) once according to the manufacturer’s protocol. The cells were washed once with MACS buffer, and passed through a cell strainer cap (Falcon; 352235). The stained cells were stored at 4 °C overnight before acquisition of MACSQuant 10 (Miltenyi). For all flow cytometry, the tubes were labeled only with mouse ID to remove any bias.

### Antibodies and dextramers

APC-conjugated B27-KK10-dextramer was prepared as described (Bethune et al., 2017; Joglekar et al., 2018). Dextramer was diluted 50-fold in MACS buffer before staining. Two panels of antibodies were used to analyze human cell engraftment as follows. Panel 1: Anti-hCD45-Pacific Blue (clone HI30; Biolegend; 304029), Anti-mCD45-FITC (clone 30-F11, Biolegend; 103108), Anti-hCD19-PE-Cy7 (clone SJ25C1, Biolegend; 363012), Anti-hCD3-APC-Cy7 (clone UCHT1; Biolegend; 300426), Anti-hCD271 (NGFR) PE (clone ME20.4; Biolegend; 345106), and Dextramer-APC. Panel2: Anti-hCD45-Pacific Blue (clone HI30; Biolegend; 304029), Anti-hCD3-APC-Cy7 (clone UCHT1; Biolegend; 300426), Anti-hCD4-PE-Cy7 (clone RPA-T4; Biolegend; 300512), Anti-hCD8-FITC (clone SK1; Biolegend; 344704), Anti-hCD271 (NGFR) PE (clone ME20.4; Biolegend; 345106), and Dextramer-APC. The gating strategies used for these panels are depicted in Fig S1 (A for Panel1 and Ds for Panel2). For IFNγ secretion assay, cells were stained with Anti-hCD271 (NGFR) PE (clone ME20.4; Biolegend; 345106), and Anti-IFNγ-BV421 (clone 4S.B3, Biolegend, 502531).

### Ex vivo selection and expansion of CD45+ cells

For ex vivo expansion of human T cells from engrafted mice, spleens were collected from euthanized mice and homogenized using the plunger of a 10 ml syringe on a 40 micron nylon filter. The filters were washed with MACS buffer (DPBS with 2% FBS) to collect cells. Spleens from 2-3 mice in each arm were pooled for selection. Human CD45+ cells were enriched using the EasySep™ Mouse/Human Chimera Isolation Kit (Stemcell technologies, 19849) according to the manufacturer’s instructions. The cells were cultured in R10 supplemented with 40 U/ml rhIL2 and Immunocult anti-CD3/28 activator for 7 days. The cells were harvested and used as effector cells in cytotoxicity assays.

### HIV infection of mice

Replication competent HIV^NL4-3^ was packaged as described below. Engrafted mice were first anesthetized under isoflurane and then tattooed on their right leg with green ink (Ketchum Inc.) to indicate infection. Mice were injected intraperitoneally with 500 ng p24 of the virus and returned to their cage. Following infection, the mice were handled with long forceps with soft tips. Any procedures after infection were performed in a dedicated BSL2+ facility.

### Cytotoxicity and cytokine secretion assays

Cytotoxicity assays using peptide-pulsed or HIV-infected cells were performed as described previously (Joglekar et al., 2018). When indicated, GXR-B27 cells were labeled with CFSE (Biolegend, 423801), pulsed with 100 μg/ml of KK10 peptide (Thermo Pierce Scientific) for 2 hours at 37 °C, and co-incubated with effector cells. When indicated, GXR-B27 cells were infected with HIV^NL4-3^ for 3 days, harvested, and co-incubated with effector cells. To measure cell survival, co-incubation reactions of effector and target cells were stained with propidium iodide (PI, Life Technologies, P3566), and acquired on MACSQuant 10. Live target cells were gated as shown in Fig S3. Target cell survival was measured by counting the CFSE+PI- or GFP+PI-cells at 24 hours after co-incubation. Target cell survival was normalized to Mock-transduced effectors as described previously. For cytokine secretion assays, activated and transduced T cells were incubated with GXR-B27 cells pulsed with the indicated peptide concentrations for 6 hours in presence of Brefeldin A (NEB; 420601). The cells were stained with anti-LNGFR-PE, fixed (Biolegend; 420801), permeabilized (Biolegend; 421002), and stained with anti-IFNγ-BV421 (clone 4S.B3, Biolegend, 502531) before acquisition on MACSQuant 10.

### Packaging replication competent HIV^NL4-3^ and titer determination

Replication competent HIV^NL4-3^ was produced by transfection of HEK-293T cells. HEK-293T cells were plated at 5×10^6^ cells on a poly-L-Lysine-coated 10 cm plate. At 24 hours, the cells were transfected with 15 μg of the pNL4-3 plasmid using BioT transfection reagent (Bioland Scientific, B01-01). The virus was harvested 72 hours later, filtered through 0.45 micron filter, and stored at −80 °C until further use. To determine the viral titer, thawed virus was lysed in Triton-X (Sigma; 9002-93-1) and used for p24 measurement at the UCLA CFAR Virology core. The infectivity of the virus was also confirmed by infecting GXR-B27 cells and measuring the frequency of GFP+ cells at 3 days by flow cytometry.

### Generation of NL4-3 mutants and confirming viral fitness

The L6M mutation was introduced in pNL4-3 using Q5 site directed mutagenesis (New England Biolabs, E0554S) using the primers 5’-ATGGATAATCatgGGATTAAATAAAATA-3’ and 5’-CTTTTATAGATTTCTCCTACTGG-3’ according to manufacturer’s instructions. Mutant plasmid was confirmed by Sanger sequencing and used to package the virus. L6M mutant virus was packaged as described above and stored at −80 °C until further use. To measure viral fitness, parallel infection of GXR-B27 cells was performed using the wildtype and L6M mutant viruses. Frequency of GFP+ cells was measured by flow cytometry at 3 days to determine infectivity.

### Viral load determination

Viral loads in infected mice were determined using a clinical droplet digital PCR assay performed at the UCSD CFAR Translational Virology core. RNA was extracted using 10 μl of stored plasma using the E.Z.N.A. viral nucleic acid isolation kit (Omega; R6874) with on-column DNase I digestion (Omega; E1091). Extracted RNA was stored at −80 °C until further use. Viral loads were measured using one-step RT-PCR based assay to measure copies of a region in the pol gene. The primers and probes used for the assay are: HIV Pol Probe: 5’-CCCACCAACAGGCGGCCTTAACTG-3’ with FAM/TAMRA; HIV-F:5’-CAATGGCAGCAATTTCACCA-3’ and HIV-R: 5’-GAATGCCAAATTCCTGCTTGA-3’. The RNA samples sent to the CFAR TV core were labeled only with mouse ID to ensure that the viral load measurement is done in a blinded fashion. The limit of detection of this assay is 0.06 copies per μl. Any samples that showed undetectable viral load were given the value of 0.05 copies per μl in order to plot them on a logarithmic scale.

### Viral sequencing

For sequencing viral quasi-species, a three-step RT-PCR approach was used. First, cDNA was generated using the Superscript IV First-Strand Synthesis System (Life Technologies; 18091050) at 55 °C for 10 minutes, 80 °C for 10 minutes, followed by RNase H digestion at 37 °C for 20 minutes. Second, the generated cDNA was subjected to a first round of PCR using Platinum SuperFi PCR Master Mix (Life Technologies; 12358250) to amplify a 3 kb region spanning gag-pol using primers Pan-HIV-1_2F: 5’-GGGAAGTGAYATAGCWGGAAC-3’ and Pan-HIV-1_2R: 5’-CTGCCATCTGTTTTCCATARTC-3’ using the following conditions: initial denaturation at 98 °C for 30 seconds, followed by 40 cycles of incubation at 98 °C for 10 seconds, 58 °C for 10 seconds, and 72 °C for 2 minutes, and a final extension of 72 °C for 3 minutes. Third, the PCR product was subjected PCR using Q5^®^ Hot Start High-Fidelity 2X Master Mix (NEB; M0494S) in two separate reactions to amplify the KK10 or the KY9 epitope. Illumina Unique Dual Indexes were incorporated into the amplicons using a 10-fold excess of the index primers. The primers and conditions used for the PCR are as follows: Primers for KK10: TruSeq-Read1-KK10-mix_1: 5’-CTCTTTCCCTACACGACGCTCTTCCGATCTGGATGGATGACACATAATCCACC-3’; TruSeq-Read1-KK10-mix_2: 5’-GTGACTGGAGTTCAGACGTGTGCTCTTCCGATCTAATGCTGGTAGGGCTATACATTC-3’; Primers for KY9: TruSeq-Read1-KY9-mix_1: 5’-CTCTTTCCCTACACGACGCTCTTCCGATCTTGAACATCTTAAGACAGCAGTACAA-3’; TruSeq-Read1-KY9-mix_2: 5’-GTGACTGGAGTTCAGACGTGTGCTCTTCCGATCTGTCTGTTGCTATTATGTCTACTATTC-3’. Primers for Indexing: i5_index: 5’-CAAGCAGAAGACGGCATACGAGA-index-GTGACTGGAGTTCAGACGTGTGCTCTTCCGATC-3’; i7_index: 5’-AATGATACGGCGACCACCGAGATCTACACAGCGCTAGACAC-index-ACACGACGCTCTTCCGATCT-3’. The PCR conditions: initial denaturation at 98 °C for 30 seconds, followed by 20 cycles of incubation at 98 °C for 10 seconds, 58 °C for 20 seconds, and 72 °C for 30 seconds; and a final extension of 72 °C for 2 minutes. The PCR products were checked by gel electrophoresis, pooled, purified using a PCR purification kit (Macherey-Nagel Nucleospin; 740609), and sent for Illumina sequencing. Paired end 150-read sequencing was performed on a HiSeq4000 (Fulgent Genetics).

### Sequence analysis

The FASTQ files obtained from the sequencer were first de-multiplexed into individual indexes. The sequencing reads were aligned to the sequence of pNL4-3 using Burrows-Wheeler Aligner. Aligned reads were translated into the epitope, and reads corresponding to each variant of the KK10 or KY9 epitope were scored. The scores for each epitope variant were reported as pie-charts using a custom script in R. Sequence analysis was performed using custom scripts in Python.

### Statistics and data analysis

Flow cytometry data were analyzed using FlowJo X (TreeStar). All statistical analyses were performed in GraphPad Prism (GraphPad Software). Plotting of the viral quasi-species was performed using R.

## Author Contributions

A.V.J. conceptualized the study, obtained funding that led to the study, designed and performed experiments, analyzed the data, and wrote the manuscript. M.S. performed in vitro and in vivo experiments, analyzed the data, and wrote the manuscript. M.T.L. performed in vitro experiments and analyzed viral sequencing data. J.D.J. and S.S. performed experiments, D.B. conceptualized the study, obtained funding that led to the study, oversaw the study, and wrote the manuscript.

## Acknowledgements

We thank the staff at Caltech’s Office of Laboratory Animal Research, particularly John Papsys, Brittney Garcia, Gwen Williams, Ruben Bayon, and Dr. Karen Lencioni for their assistance with mouse colony maintenance, timed matings, and maintenance of animal health. We thank Caroline Ignacio and Gemma Caballero at the University of California, San Diego, Center for AIDS Research Translational Virology Core for running ddPCR measurement for viral load. We thank Deborah Anisman-Posner and Alex Bollinger at the University of California, Los Angeles, Center for AIDS Research Virology Core for collecting PBMCs from healthy donors and for running p24 measurement from viral supernatant. We thank the staff at the Fred Hutchinson Cancer Research Center Core in Experimental Hematology for screening HSPC donors and for collecting and enriching mPB-HSPCs from healthy donors. We thank Rishi Bhargava for assistance with generating pie charts depicting the frequencies of viral quasi-species. We thank Dr. Anjie Zhen, Dr. Scott Kitchen, and members of the Baltimore laboratory for discussions and suggestions regarding the study. We thank Zhe Liu and Won-Jun Noh for technical assistance. Finally, we thank Dr. Bruce D. Walker at the Ragon Institute for MGH, MIT, and Harvard for providing HIV controller samples from which the TCRs were isolated, as well as for scientific insights and discussions. This study was funded by the California Institute for Regenerative Medicine grant DISC2-09123 and California Institute of Technology Innovation Initiative.

---

**Fig S1:**
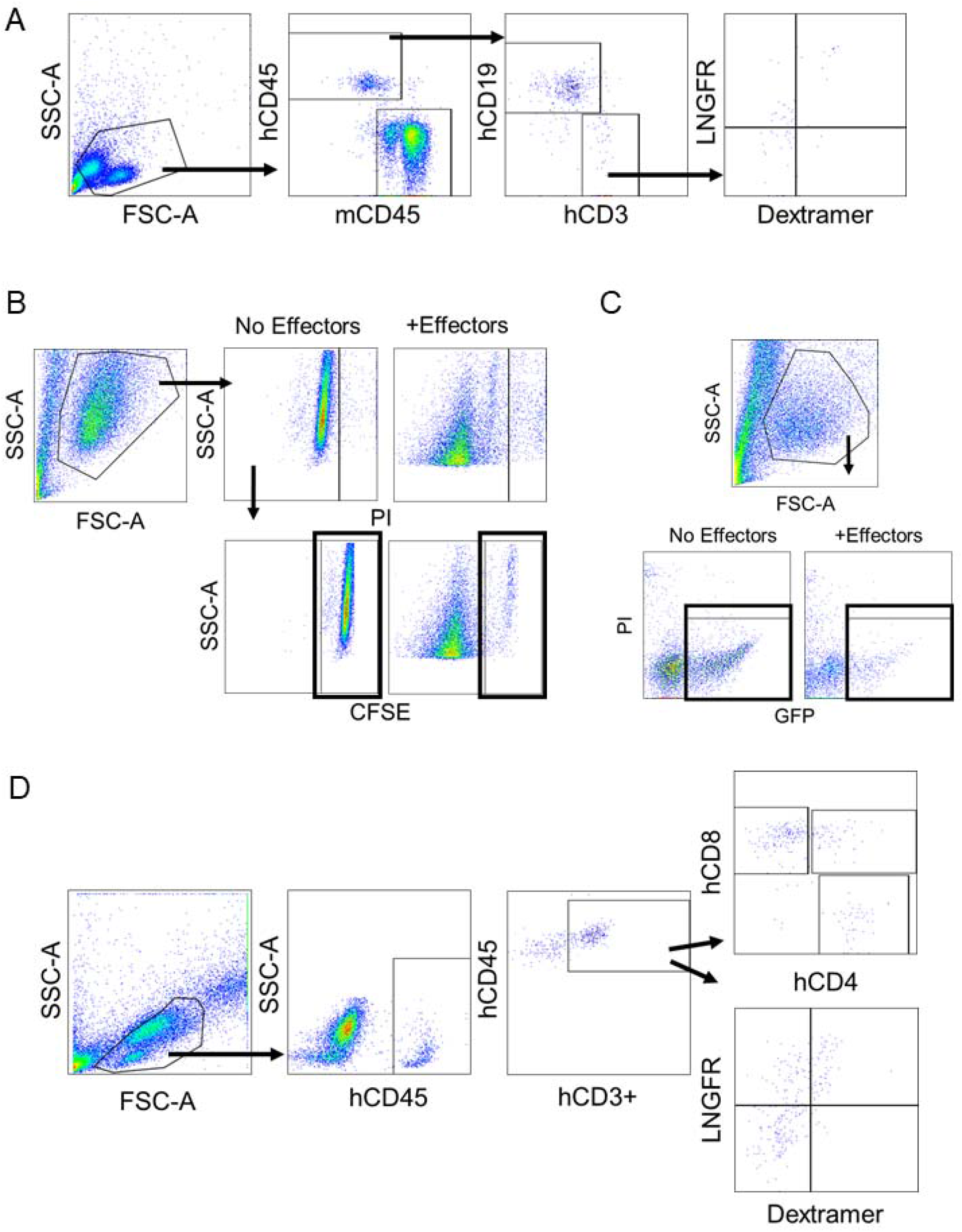
Gating strategy used for flow cytometry. Gating strategy used for A. analyzing human cell engraftment using antibody panel 1, B. analyzing cytotoxicity assays using peptide-pulsed target cells, C. analyzing cytotoxicity assays using HIV-infected target cells. D. analyzing human cell engraftment using antibody panel 2. In B and C, black rectangles indicate gates used to count live target cells.

**Fig S2:**
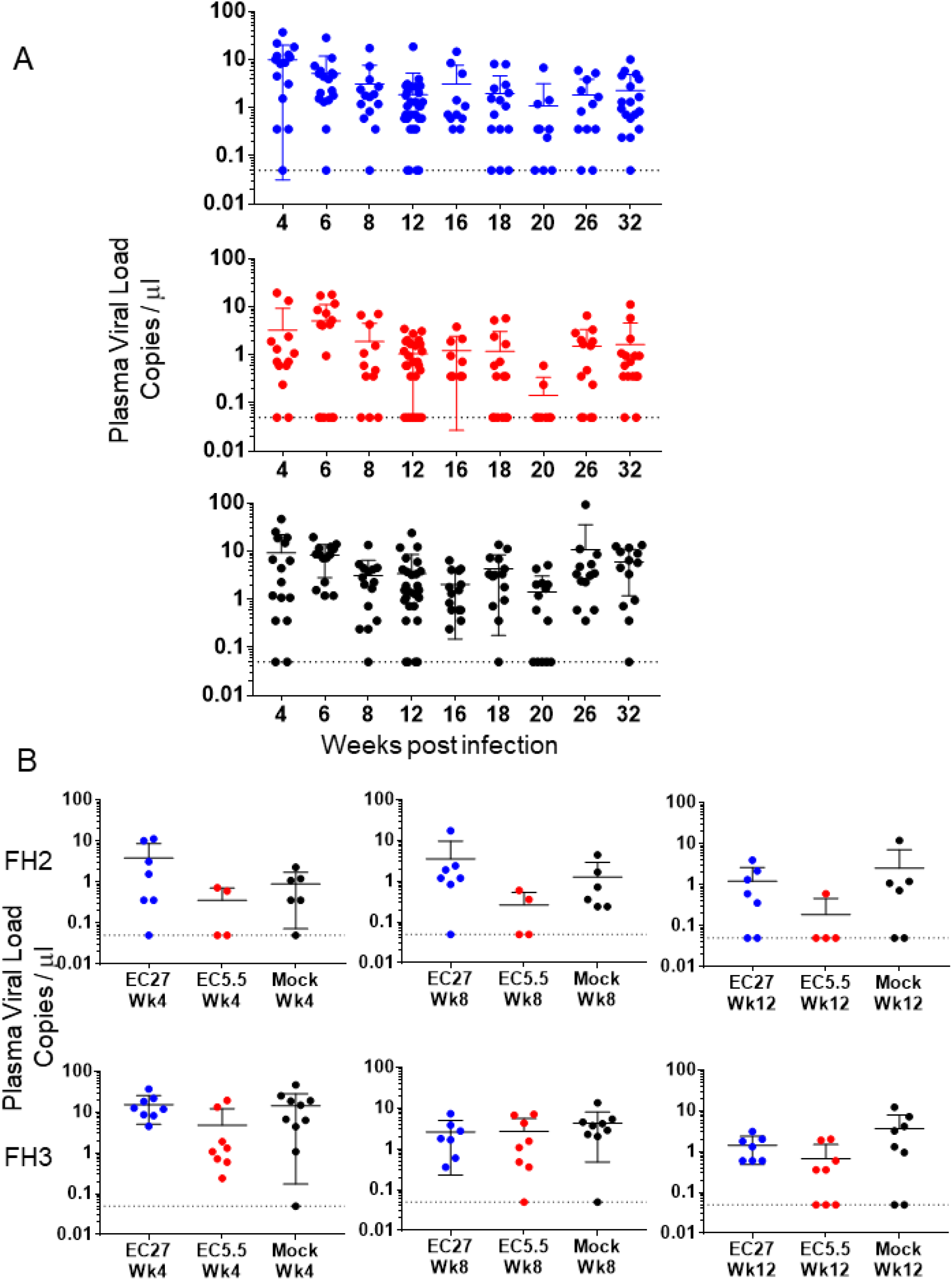
Individual data points for plasma viral load measurements. A. Viral load in the plasma from peripheral blood of HIV-challenged mice in both cohorts across all time points. Dots indicate individual mice. Mean and S.D. are indicated by solid lines and error bars. Dotted line indicates the limit of detection (0.05 copies per μl). Top graph: EC27-mice, Middle graph: EC5.5-mice, Bottom graph: Mock-mice. B. Viral load in the plasma from peripheral blood of HIV-challenged mice in cohort22-23 at 4, 8, and 12 weeks post-infection. The top and bottom graphs show data at the same time point for humanized mice generated using two different HSPC donors, FH2 and FH3 as indicated. Dots indicate individual mice. Mean and S.D. are indicated by solid lines and error bars. Dotted line indicates the limit of detection (0.05 copies per μl).

**Fig S3:**
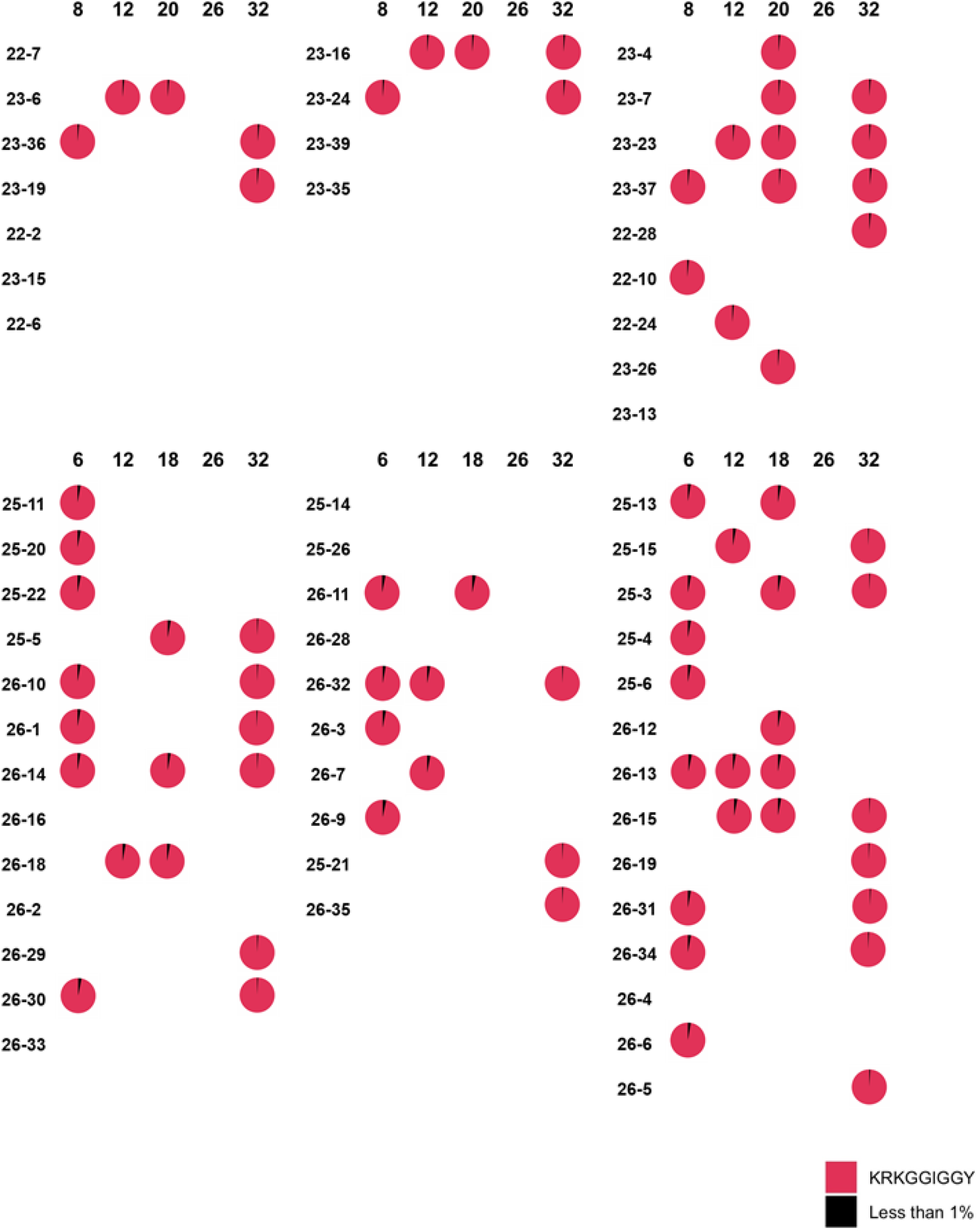
Evolution at the KY9 epitope in viral quasi-species in infected mice. Each pie chart indicates all viral quasi-species encompassing over 1% of the total sequence diversity at the KY9 epitope. For each TCR, mouse IDs are shown on the left and the week post-infection at which the sample was obtained is indicated on top. The top part of the figure shows data from cohort22-23, the bottom part shows data from cohort 25-26. The legend for the individual colors and their corresponding mutant epitopes is at the bottom.

